## Supplemental figures 1-3 and movie legends for "Deep learning for rapid analysis of cell divisions *in vivo* during epithelial morphogenesis and repair"

### Supplemental Information

*Jake Turley[1,2], Isaac V. Chenchiah[1,*], Paul Martin[2,*] , Tanniemola B. Liverpool[1,*], Helen Weavers[2,*]*

** co-corresponding authors; these authors contributed equally.*

*Institutional affiliations:*
*[1]School of Mathematics, University of Bristol, Bristol, BS8 1UG, UK*
*[2]School of Biochemistry, University of Bristol, Bristol, BS8 1TD, UK*

*Author Emails:*
**

### Supplemental Figures

Further analysis of division density in living epithelial tissue *in vivo*

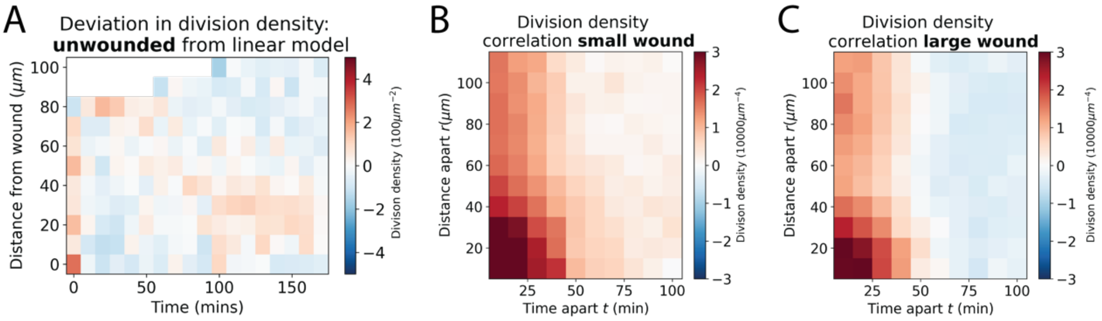


***Figure S1. Further analysis of division density in living epithelial tissue.*** *A) Heatmaps of the deviation in division density of unwounded tissue compared with a best fit linear model. The axes are time and distance from a virtual wound. Red areas have more division and blue less. B-C) Heatmaps of the division density correlation for small and large wounds. Again, red areas have more divisions and blue less. Also see Figure 3.*

UNetOrientation model diagram and error of test data

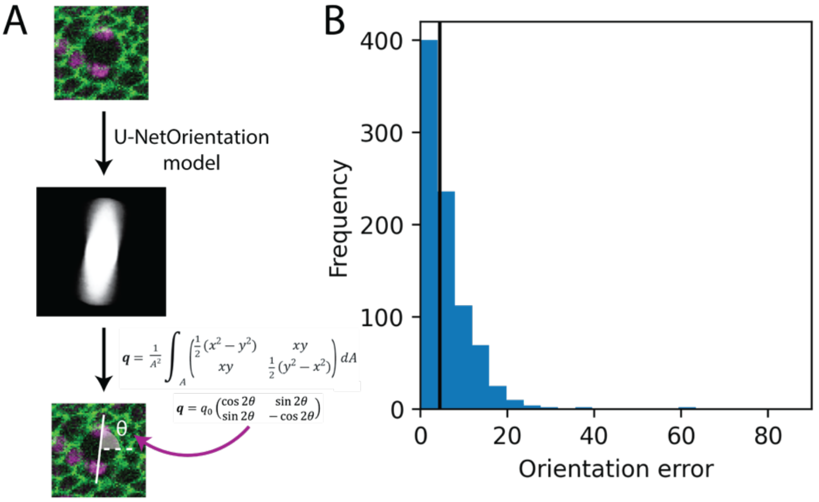


***Figure S2**. **UNetOrientation model diagram and error of test data**. A) Diagram of the output of U-NetOrientation. The oval is elongated in the same direction as the division, thus calculating its q-tensor tells us the orientation of the cell division. B) The error of the U-NetOrientation model on the test dataset; black line shows median error. Also see Figure 4.*

### Further analysis of division orientation in living epithelial tissue *in vivo*

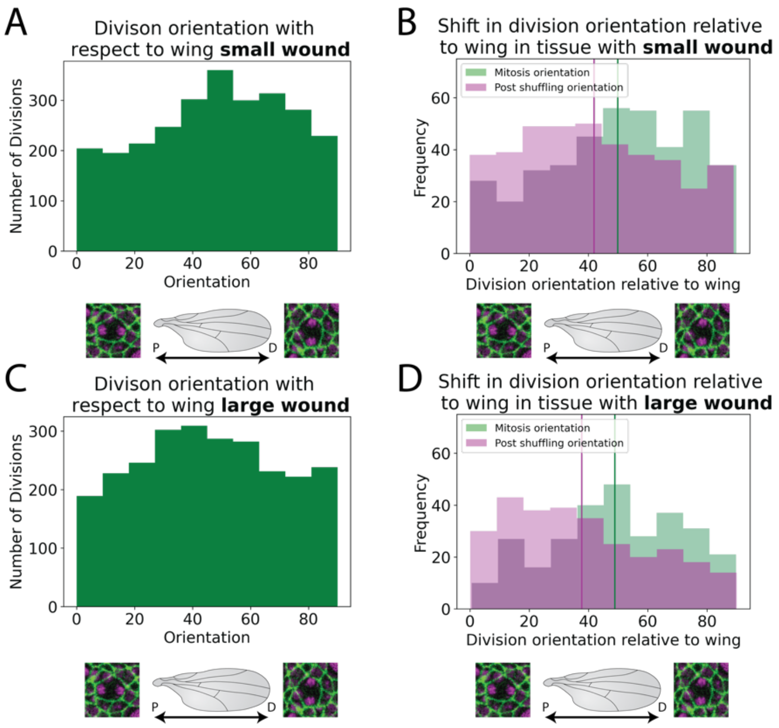


***Figure S3. Further analysis of division orientation in living epithelial tissue.*** *A) Distribution of the division orientations for small wounds. B) Distribution of the division orientations with respect to the wing in small wounds (green), and the daughter cell orientation 20 mins after dividing (magenta). C) Distribution of the division orientations for large wounds. D) Distribution of the division orientations with respect to the wing in large wounds (green), and the daughter cell orientation 20 mins after dividing (magenta). Also see Figure 4.*

### Supplemental Movie Legends

**Movie S1. Time-lapse imaging of the unwounded pupal epithelium over 3 hours.** Projected from a 3D stack using the stack focus algorithm with a radius of 5 pixels. Green indicates *E-cadherin-GFP* and magenta indicates *Histone2-RFP*. The white circles show the divisions detected by the "U-NetCellDivision10" and the white lines indicate the orientation of divisions determined by "U-NetOrientation". Scale bar: 10 μm. Related to Figure 3.

**Movie S2. Time-lapse imaging of a small wound in the pupal epithelium over 3 hours.** Projected from a 3D stack using the stack focus algorithm with a radius of 5 pixels. Green indicates *E-cadherin-GFP* and magenta indicates *Histone2-RFP*. The white circles show the divisions detected by the "U-NetCellDivision10" and the white lines indicate the orientation of divisions determined by "U-NetOrientation". Scale bar: 10 μm. Related to Figure 3.

**Movie S3. Time-lapse imaging of a large wound in the pupal epithelium over 3 hours.** Projected from a 3D stack using the stack focus algorithm with a radius of 5 pixels. Green indicates *E-cadherin-GFP* and magenta indicates *Histone2-RFP*. The white circles show the divisions detected by the "U-NetCellDivision10" and the white lines indicate the orientation of divisions determined by "U-NetOrientation". Scale bar: 10 μm. Related to Figure 3.

**Movie S4. Time-lapse imaging of a small wound in the pupal epithelium over 3 hours.** Projected from a 3D stack using the stack focus algorithm with a radius of 5 pixels. Grayscale background of epithelium with circles show the divisions detected by the "U-NetCellDivision10", the lines indicate the orientation of divisions determined by "U-NetOrientation" and the colour of labels display the orientations relative to wounds. Blue labelled divisions are orientated towards wounds, red away from wounds and white around 45°. The white dot is the centre of the wound and the closed wound site after closure. Scale bar: 10 μm. Related to Figure 3.

**Movie S5. Time-lapse imaging of a large wound in the pupal epithelium over 3 hours.** Projected from a 3D stack using the stack focus algorithm with a radius of 5 pixels. Grayscale background of epithelium with circles show the divisions detected by the "U-NetCellDivision10", the lines indicate the orientation of divisions determined by "U-NetOrientation" and the colour of labels display the orientations relative to wounds. Blue labelled divisions are orientated towards wounds, red away from wounds and white around 45°. The white dot is the centre of the wound and the closed wound site after closure. Scale bar: 10 μm. Related to Figure 3.

### Supplemental Tables

**Table 1 : Dice scores for the deep learning models**

| Model | True Positives | False Positive | False Negative | Dice score |
|---|---|---|---|---|
| U-NetCellDivision3 | 797 | 216 | 310 | 0.752 |
| U-NetCellDivision10 | 1057 | 28 | 50 | 0.964 |

**Table 2 : Dice scores for the segmentation methods**

| Segmentation | True Positives | False Positive | False Negative | Dice score |
|---|---|---|---|---|
| Single focal plane + Tissue Analyzer | 8,197 | 313 | 4,317 | 0.780 |
| U-NetBoundary + Tissue Analyzer | 11,325 | 501 | 1,189 | 0.931 |
